# The H3K4me3 binding ALFIN-LIKE proteins recruit SWR1 for gene-body deposition of H2A.Z

**DOI:** 10.1101/2024.09.12.612642

**Authors:** Linhao Xu, Yafei Wang, Xueying Li, Qin Hu, Vanda Adamkova, Junjie Xu, C. Jake Harris, Israel Ausin

## Abstract

The H2A.Z histone variant is highly enriched over gene bodies, playing an essential role in several genome-templated processes, including transcriptional regulation and epigenetic patterning across eukaryotes. The SWR1 chromatin remodeling complex deposits H2A.Z. How SWR1 is directed to gene bodies is largely unknown. Here, we show that ALFIN-LIKE (AL) proteins are responsible for H2A.Z gene body patterning in *Arabidopsis*. AL proteins encode H3K4me3-binding PHD domains, and by ChIP-seq, we confirm preferential binding of AL5 to H3K4me3 over H3K4me1/2 *in planta*. We observe a global reduction in H2A.Z in *al* septuple mutants (*al7m*), especially of over H3K4me3-enriched genic regions. While MBD9 recruits SWR1 to nucleosome-free regions, ALs act non-redundantly with MBD9 for deposition of H2A.Z. Notably, *al7m* mutants show severe developmental abnormalities and upregulation of H2A.Z gene body-enriched responsive genes. Therefore, we propose a model whereby AL proteins direct gene body enrichment of H2A.Z by recruiting SWR1 to H3K4me3-containing responsive genes.

## Introduction

A unique feature of eukaryotic genomes is that their genetic material is organized into chromatin. During chromatin formation, approximately 147 DNA base pairs wrap around a histone octamer consisting of a central H3-H4 tetramer and one H2A-H2B dimer on each side. The resulting structure - known as the nucleosome - serves as the fundamental repeating unit of chromatin (1–3). While chromatin affords opportunities for genome organization and packaging, nucleosomes represent an inherent obstacle to cellular factors that require access to DNA during processes such as replication, transcription, and DNA repair. Therefore, after initial assembly, chromatin often undergoes dynamic changes in composition, density, and/or post-translational modifications to ensure accessibility and function (4).

One of these dynamic changes in composition involves the exchange of canonical histones by their variants. Histone variants are paralogs that differ from their canonical partners by a few amino acids or short specific domains (5, 6). The histone variant H2A.Z is present in all eukaryotic lineages from yeast to metazoans. It has been shown to play crucial roles in several biological processes, such as DNA repair, recombination, or transcription (6, 7). Interestingly, H2A.Z is associated with both transcriptional stimulation and repression in different genomic contexts, although the molecular mechanisms remain unclear. H2A.Z is generally present at the first nucleosome immediately downstream of the transcriptional start site, i.e., the +1 nucleosome. However, H2A.Z occupancy steeply declines in highly expressed genes in subsequent nucleosomes. In contrast, low-expressed genes are associated with high H2A.Z levels across the gene body (8–10). H2A.Z can also undergo post-translational modifications that affect transcription. For instance, ubiquitination of H2A.Z can result in repression, while acetylation of H2A.Z may promote activation (8). Genome-wide, H2A.Z is predominantly located in euchromatic regions and is absent from constitutive heterochromatin (11, 12). However, it is also present in facultative heterochromatin regions along with H3K4me3, H3K27me3, and H2A-Ub (13, 14).

H2A.Z is incorporated into chromatin by the chromatin remodeler SWI/SNF-Related1 complex (SWR1), which replaces H2A-H2B with H2A.Z-H2B dimers (15–17). Recent studies have used genetic and biochemical approaches to identify the components of the Arabidopsis SWR1. Most of the SWR1 complex components were found to be conserved from yeast, but additional partners were discovered, including METHYL BINDING DOMAIN9 (MBD9), four members of the ALFIN-LIKE family (AL4, 5, 6, and 7), and two subunits of the SAGA complex, TRANSCRIPTION-ASSOCIATED PROTEIN 1A and 1B (18–20).

How the SWR1 complex is recruited to chromatin remains to be fully resolved. Studies on Arabidopsis and yeast indicate that H3/H4 acetylation may play an important role as the H4 acetyltransferase complex, Nua4, is necessary to maintain H2A.Z patterning, particularly around the +1 nucleosome (21–26). Furthermore, HISTONE DEACETYLASE 9 (HDA9) and POWERDRESS are required for H2A.Z removal at the *YUCCA8* promoter during thermomorphogenesis (27, 28). Besides histone acetylation, MBD9 may also direct SWR1 to the promoter of actively transcribed genes (19, 29)

In Arabidopsis, ALs constitute a family of seven members (AL1 to 7) that share high levels of similarity(30–32). Recombinant AL proteins from Arabidopsis can bind H3K4me2/3 peptides through their PHD domains (30, 32). *ALFINs* were initially identified as genes involved in *Medicago sativa’s* salt-tolerance response (33), and the Arabidopsis ALs have also been implicated in various abiotic stress and physiological processes, such as phosphate starvation, drought, osmotic stress, cold tolerance (34–38), germination, skotomorphogenesis (37, 39) as well as epigenetic silencing (40). However, a precise molecular mechanism for the function of AL proteins remains elusive.

Here, we show that the absence of functional AL proteins leads to abnormal development and misregulation of genes bearing the hallmarks of facultative heterochromatin. We show that AL5 interacts with both H3K4me3 and the SWR1 complex *in vivo*. Importantly, *al* mutants show a significant reduction in H2A.Z at loci that coincide with AL5 binding. Our findings suggest that ALs recruit the SWR1 complex to chromatin via binding to H3K4me3, facilitating the deposition of H2A.Z at stress-responsive loci.

## Results

### AL proteins are required for proper development and the control of inducible genes

The seven Arabidopsis ALs are highly related (30–32, 39), and the *al* single mutants do not show any obvious developmental phenotypes under standard growing conditions (38, 39) (Supplemental Figure 1C). To overcome any potential functional redundancy, we generated higher-order mutants. We obtained *al1-2* to *al3-1* and *al5-1* to *al7-1* from a publicly available seed bank (https://abrc.osu.edu) (Supplemental Figure 1A, 1B). However, we could not obtain an *AL4* mutant allele; thus, we generated an *al4-2* allele using CRISPR (Supplemental Figure 1A, 1B). The final homozygous AL septuple mutant *al1-2 2-1 3-1 4-2 5-1 6-1 7-1* (hereafter referred to as ‘*al7m’*) shows dramatic developmental defects, including low viability, slow growth, dwarfism, curled leaves, reduced apical dominance, abnormal flowers, and sterility (Figure 1A, Supplemental Figure 1D), indicating that ALs play a significant role in development.

**Figure 1.**
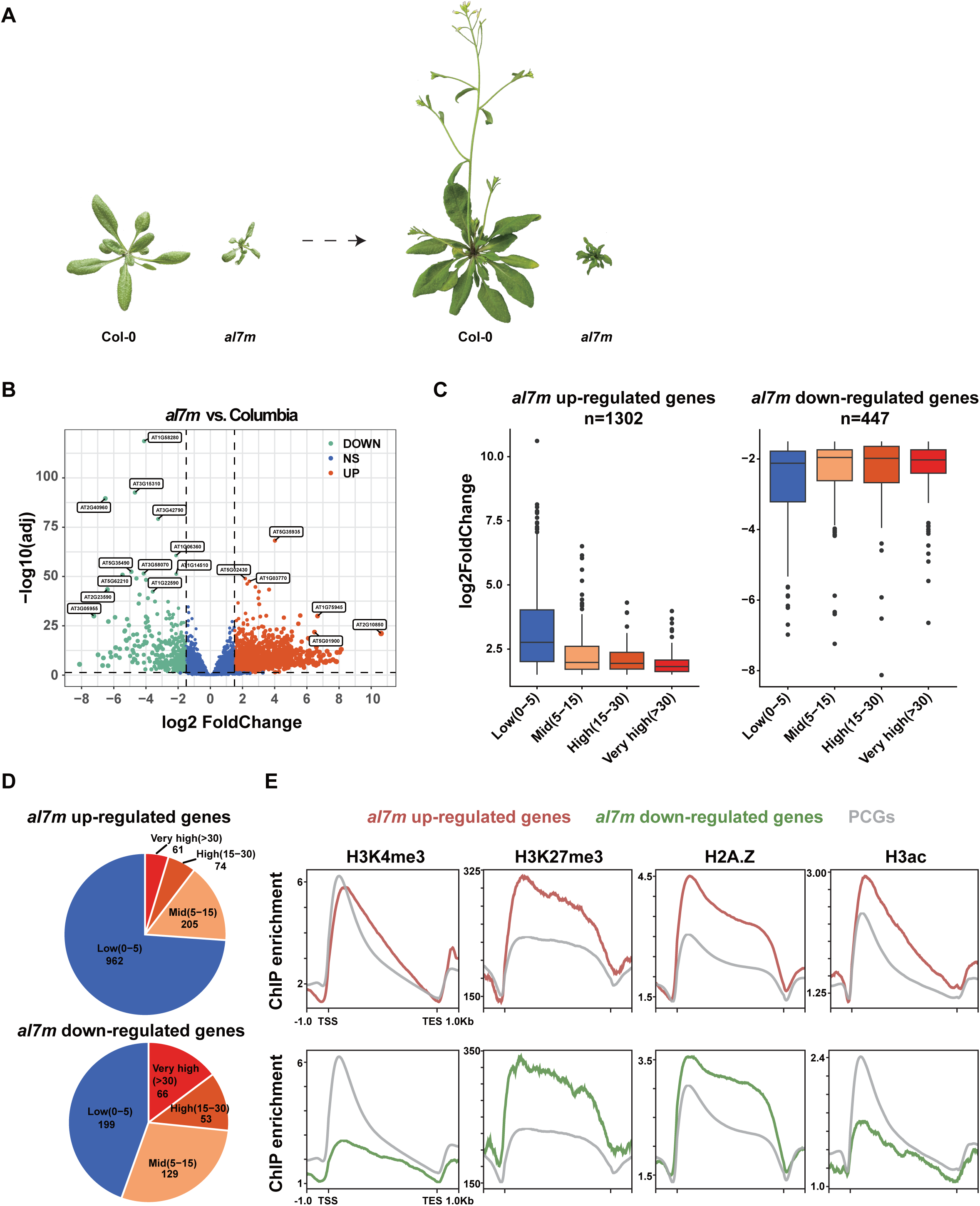
alfin-like septuple mutant (al7m) phenotype and analysis of the al7m transcriptome. (A) The visual phenotype of the wildtype and al7m on 28 (left) and 40 (right) days after germination. (B) Volcano plot showing deregulated genes in the al7m mutant background. A cut-off of log2>±1.5 and FDR<0.05 was used for the differential expression level and statistical significance, respectively. (C) Box plots showing differential expression of deregulated genes in al7m divided by expression level classes in the wild type. The boxplot’s top, mid-line, and bottom represent the upper quartile, median, and lower quartile, respectively. (D) Pie charts showing the number of deregulated genes in al7m separated according to gene expression categories in the wild type. (E) Metaplots showing the typical profile of the epigenetic marks H3K4me3, H3K27me3, H2A.Z, and H3ac over deregulated genes in the al7m background and all protein-coding genes (PCGs).

AL proteins have also been implicated in the response to several abiotic stresses (34–38), suggesting that their role is not restricted to development. To better understand AL function, we performed RNA-seq in the *al7m* and the wild-type control (WT, Col-0). This revealed 1749 genes that are significantly affected by the lack of functional ALs. Among these, approximately three-quarters (1302) are upregulated, while the rest are downregulated (Figure 1B). Most upregulated genes (960) typically have low expression in WT (Figure 1C and 1D). In contrast, approximately half of the downregulated are low expressed in WT (199), and the remaining sit within the mid (129), high (53), and very high (66) expression categories (Figures 1C and 1D). Gene ontology analysis indicates that upregulated genes are highly enriched for responsive genes (Supplemental Figure 2). Together, these data indicate that the ALs primarily act by repressing transcriptional activity at inducible genes under conditions where its expression is not required.

Next, we checked some key epigenetic features of the genes deregulated in the *al7m* background. H3K27me3 levels and H2A.Z occupancy were higher over the *al7m* deregulated genes than the overall average of protein-coding genes, consistent with these deregulated genes being low expressed in WT (Figure 1E). However, the upregulated genes in the *al7m* background also showed surprisingly high H3K4me3 and H3K9/14Ac levels, which are two epigenetic marks typically associated with high transcriptional activity (Figure 1E). These data suggest that AL proteins may specialize in repressing low-expressed genes with a bivalent chromatin signature.

### AL proteins bind H3K4me3 in planta

The AL proteins have been shown to bind to H3K4me3 *in vitro* by pull-down with modified histone peptides. AL1,3,4,5 and 7 can also bind to H3K4me2 peptides with lower affinity (30, 32). To investigate AL binding *in vivo*, we generated a complementing line (Supplemental Figure 3) expressing AL5 fused to nine copies of the Myc epitope. Performing co-immunoprecipitation (Co-IP) analyses, we confirmed that AL5 physically interacts with H3K4me3 containing chromatin *in planta* (Figure 2A).

**Figure 2.**
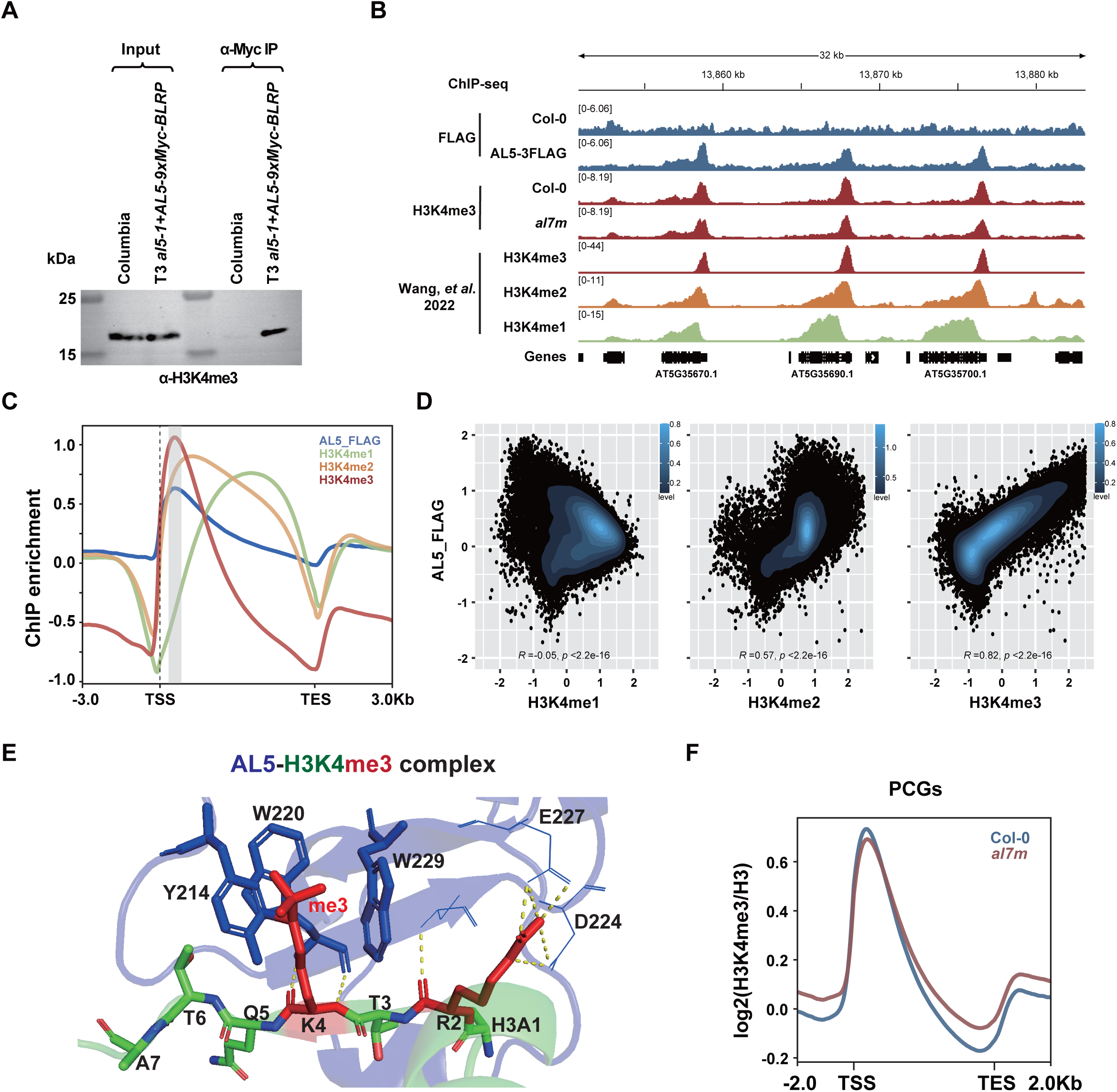
AL5 is an H3K4me3 binder. (A) Co-immunopurification assays confirming the interaction of AL5 with H3K4me3 in vivo. Immunoprecipitation lane (α-Myc IP) showing co-purification of AL5. The antibody used to detect the target is indicated at the bottom of the blot. Extracts from the same lines are included to identify the co-precipitating band (Input). (B) Genome browser image showing the colocalization of AL5-FLAG peaks with H3K4me3 peaks at selected genes. (C) Metaplot showing the ChIP-seq signals of AL5-FLAG and published data sets of H3K4me1, H3K4me2 and H3K4me3 (Wang, et al. 2022) over all genes. (D) Scatterplots showing the correlation of AL5-FLAG and H3K4me (Wang, et al. 2022) over all PCGs. Spearman was used as a correlation method. The colored bin represents the density level of dots. (E) AlphaFold three predicted the structure models of the AL5-H3K4me3 complex. The AL5 is shown in blue, H3 is shown in green, and the histone H3K4me3 and H3R2 peptides are shown in red. AL5 residues involved in H3K4me3 recognition are labeled. (F) Metaplot showing the ChIP-seq signal of H3K4me3 in Col-0 and al7m over PCGs.

Next, we performed ChIP using the *AL5-3xFlag* complementing line. (Supplemental Figures 3 and 4A to C). Comparison with available data (41) showed a striking resemblance of the general profile of AL5 to that of H3K4me3 occupancy (Figure 2B). Metaplot profiles over protein-coding genes showed that AL5 enrichment peaks directly over that of H3K4me3, while H3K4me2 and H3K4me1 occupancies peak further downstream in the gene body (Figure 2C). To quantify the correspondence between AL5 and H3K4me1/2/3, we performed linear regression, revealing a strong positive correlation with H3K4me3 (R^2^=0.82), and reduced associations with H3K4me2 (R^2^=0.57) and H3K4me1 (R^2^=-0.05) (Figure 2D). We also predicted the structure of the AL5-H3K4me3 complex using AlphaFold 3 (42). The aromatic cage, formed by residues Y214, W220, and W229, along with the H3R2 pocket created by D224 and E227, ensures the binding of AL5 to H3K4me3 (Figure 2E, Supplemental Figure 5). These results resemble the solved structure of the AL1-H3K4me3 complex (43). Furthermore, the AL5 signal positively correlates with gene expression (Supplemental Figure 4D), which is a well-known characteristic of H3K4me3. Together, the data indicate that AL5 is preferentially recruited to H3K4me3 sites *in vivo*.

Next, we performed H3K4me3 ChIP-seq in WT vs *al7m* to assess whether ALs are required for H3K4me3 stability (Supplemental Figure 6). We observed minimal perturbation over all genes in aggregate and both increased and decreased levels of H3K4me3 at genes that are up-regulated or down-regulated in the *al7m*, respectively (Figure 2F, Supplemental Figure 7). This correspondence indicates that any changes to H3K4me3 levels in the *al7m* are likely secondary consequences of altered gene expression. To further assess AL’s impact on H3K4me3, we focused on AL5 binding sites, finding that fewer than 6% of AL5-bound loci experience loss of H3K4me3 in the *al7m* (876 out of 14866, Supplemental Figure 6D and 8B). ALs, therefore, do not appear essential for H3K4me3 maintenance and likely act as true ‘readers’ of H3K4me3 with downstream functions.

### AL proteins physically interact with the SWR1 complex in planta

Previous studies have reported a physical interaction between the SWR1 complex and several AL proteins (18–20). We performed co-immunoprecipitation assays (Co-IP) between AL5 and the SWR1 complex core subunit ARP6, confirming a physical interaction between these components *in vivo* (Figure 3A). We also performed immunoprecipitation followed by mass-spectrometry (IP-MS) of AL5-9xMyc (Supplemental Figure 3) and identified several components of the SWR1 complex (Supplemental Table 1). These data confirm that AL proteins interact with the SWR1 complex *in planta*.

**Figure 3.**
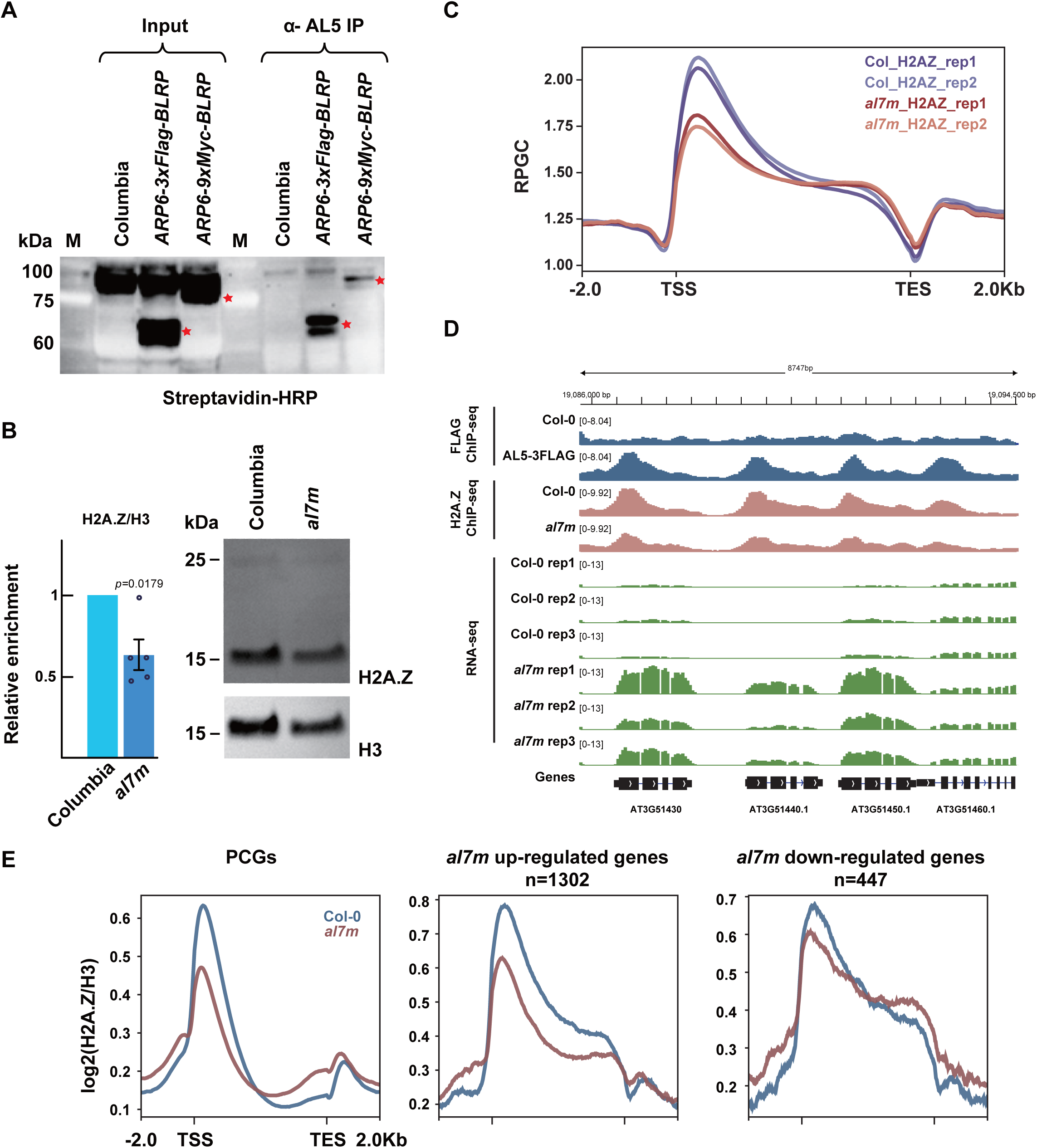
ALs are required for the deposition of H2A.Z. (A) Co-immunopurification assays confirming the interaction of AL5 with ARP6 in vivo. Immunoprecipitation lane (α-AL5 IP) showing co-purification of AL5. The antibody used to detect the target is indicated at the bottom of the blot. Extracts from the same lines are included to identify the co-precipitating band (Input). The red star indicates ARP6-3xFLAG-BLRP or ARP6-9xMYC-BLRP bands in both input and IP samples. (B) H2A.Z abundance in al7m. Western blot (right) and quantification of H2A.Z relative to H3 levels (left) in total histone extracts from the wild type and al7m. The upper band in the H2A.Z blot likely represents the ubiquitinated form of H2A.Z. Only the lower band was used for quantification purposes. The data are presented as the average levels of five independent replicates ± SE. A paired two-tailed Student’s t-test was used to determine the significance between wildtype and al7m. (C) Metaplot showing the normalized ChIP-seq signal (RPGC) of H2A.Z in two biological replicates of Col-0 and al7m over all PCGs. (D) Genome browser image showing the FLAG ChIP-seq signals of AL5-3FLAG, H2A.Z ChIP-seq signals of Col-0 and al7m, as well as the RNA-seq signals of three biological replicates of Col-0 and al7m over representative al7m up-regulated genes. (E) Metaplots showing the ChIP-seq signal of H2A.Z in Col-0 and al7m over PCGs and al7m deregulated genes.

### AL proteins are required for the deposition of H2A.Z

As the SWR1 complex is responsible for depositing H2A.Z, we reasoned that ALs might be required for H2A.Z chromatin incorporation. By western blot, we observed an approximately 40% reduction in bulk H2A.Z levels in the *al7m* as compared to WT (Figure 3B). Next, we conducted ChIP-seq analyses of H2A.Z in WT vs *al7m*. Consistent with the western blot data, this revealed a clear reduction of H2A.Z signal in *al7m* over protein-coding genes. Loss of H2A.Z was most prominent at the 5’ end of the transcriptional unit, consistent with the pattern of maximal AL5 enrichment over genes (Figure 3C to 3E). Comparing H2A.Z to H3K4me3, we observed a major reduction of H2A.Z, but not H3K4me3, directly over AL5 binding loci (Figure 4A) in the *al7m* background. Calling peaks, we found that approximately 41% of AL5 peaks experience coincident loss of H2A.Z in *al7m* (6095 out of 14866) (Supplemental Figure 6B and 8A). To determine whether the loss of H2A.Z is a general effect or is specific to regions bound by AL, we directly compared the loss of H2A.Z over AL5-bound vs. unbound loci, finding a significantly stronger reduction over AL5-bound loci (Figure 4B). Finally, we compared the dynamics of H2A.Z occupancy to the enrichment of AL5 over the transcriptional unit, finding a clear anticorrelation, whereby regions most enriched for AL5 experienced the largest losses of H2A.Z in *al7m*. Interestingly, the *al7m* upregulated genes experience loss of H2A.Z throughout the gene body, whereas downregulated genes more closely resembled the pattern over all protein-coding genes, consistent with the downregulated geneset containing both direct and indirect AL5 targets (Figure 4C). Together, these data support a model whereby ALs are required for the maintenance of H2A.Z over H3K4me3 regions bound by AL5 (Figure 6).

**Figure 4.**
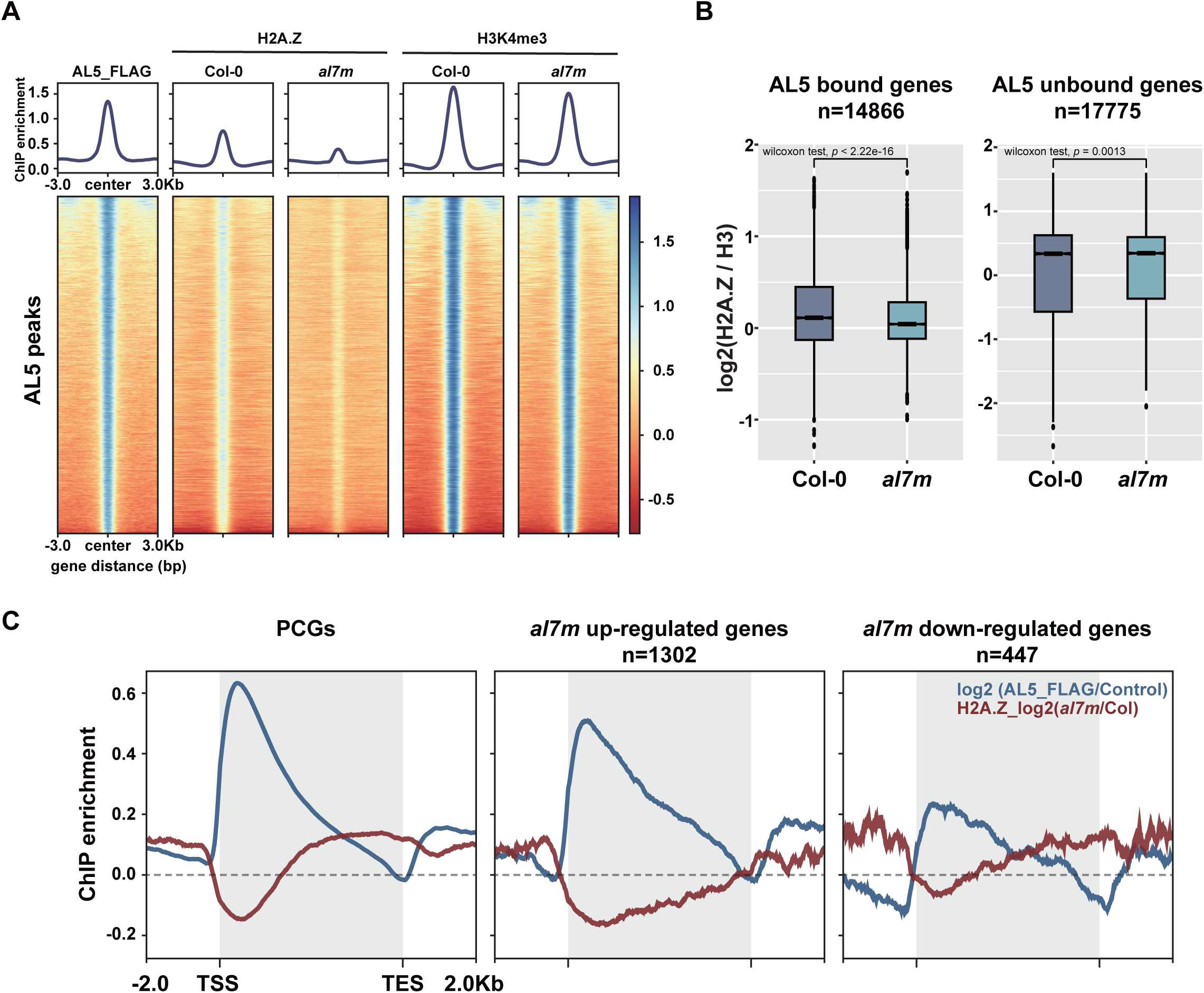
ALs repress transcription via gene body H2A.Z deposition. (A) Metaplots and heatmaps showing the ChIP-seq signals of FLAG, H2A.Z, and H3K4me3 over AL5 peaks (n=15705) in Col-0 and al7m. (B) Boxplots showing the H2A.Z levels over AL5 bound genes (n=14866) and AL5 unbound genes (n=17775). The boxplot’s top, mid-line, and bottom represent the upper quartile, median, and lower quartile, respectively. Wilcoxon test was used to determine the significance. (C) Metaplots showing the ChIP-seq enrichments of AL5-FLAG and H2A.Z over all PCGs and al7m deregulated genes.

As we previously identified MBD9 as a key component of SWR1 recruitment and H2A.Z deposition, we were interested in comparing the binding pattern of AL5 to MBD9. Using MBD9 ChIP-seq data (19) we observed that MDB9 peaks directly upstream of the transcriptional start site (TSS, as previously reported) while AL5 peaks, downstream of the TSS (Figure 5A). To gain a higher resolution perspective on the role of nucleosome positioning, we mapped the AL5 and MBD9 ChIP-seq data over the TSS in comparison to MNase-seq data (44). As previously reported, maximal MBD9 enrichment corresponded to the nucleosome-free region directly upstream of the TSS. Interestingly, AL5 peaked over the +2 nucleosome downstream of the TSS (Figure 5B). As both MBD9 and AL5 appear involved in H2A.Z deposition but show some distinct patterns of localization throughout the genome, we asked whether they function redundantly or independently. We identified 7004 MBD9 and 10954 AL5 unique peaks and 4751 peaks that were common. Over the MBD9 unique peaks, H2A.Z was significantly depleted in *mbd9-3* but not in *al7m*, while we observed the reciprocal at AL5 unique peaks. Importantly, at the common peaks, both *mbd9-3* and *al7m* were required for H2A.Z maintenance (Figure 5C). The data show that MBD9 and ALs function non-redundantly at partially overlapping regions of the genome to maintain H2A.Z patterning.

**Figure 5.**
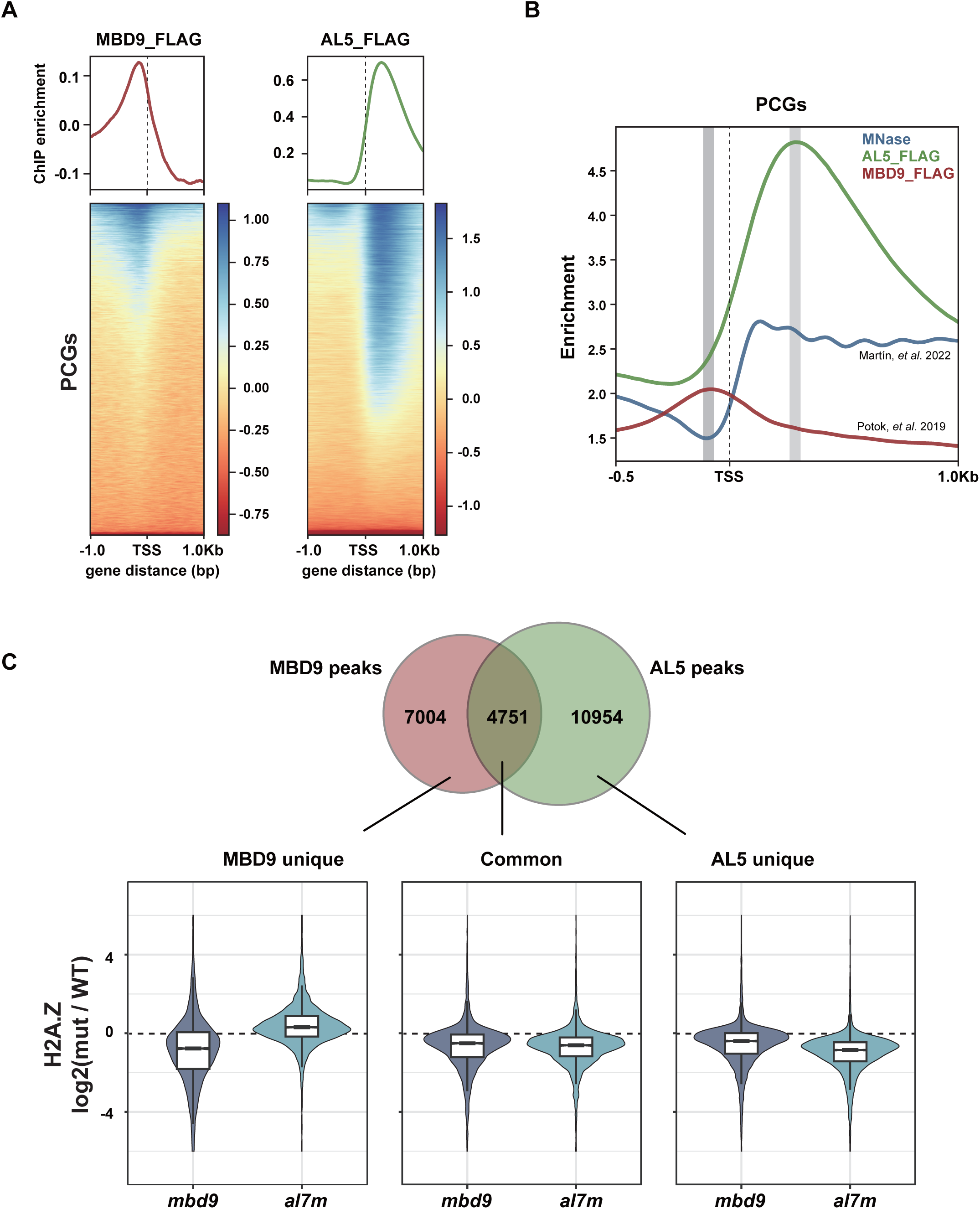
The comparison of AL5-FLAG and MBD9-FLAG ChIP-seq. (A) Metaplots and heatmaps showing the FLAG ChIP-seq signals of MBD9-FLAG (Potok, et al. 2019) and AL5-FLAG over all PCGs. (B) Metaplot showing the binding position of MBD9-FLAG (Potok, et al. 2019), AL5-FLAG and the position of nucleosomes around TSS from MNase-seq (Martín, et al. 2022). (C) Venn diagram (top) analysis of MBD9 peaks and AL5 peaks, and violin plot inlaid with boxplot (bottom) showing H2A.Z levels over MBD9 unique peaks (n=7004), common peaks (n=4751) and AL5 unique peaks (10954). The boxplot’s top, mid-line, and bottom represent the upper quartile, median, and lower quartile, respectively.

**Figure 6.**
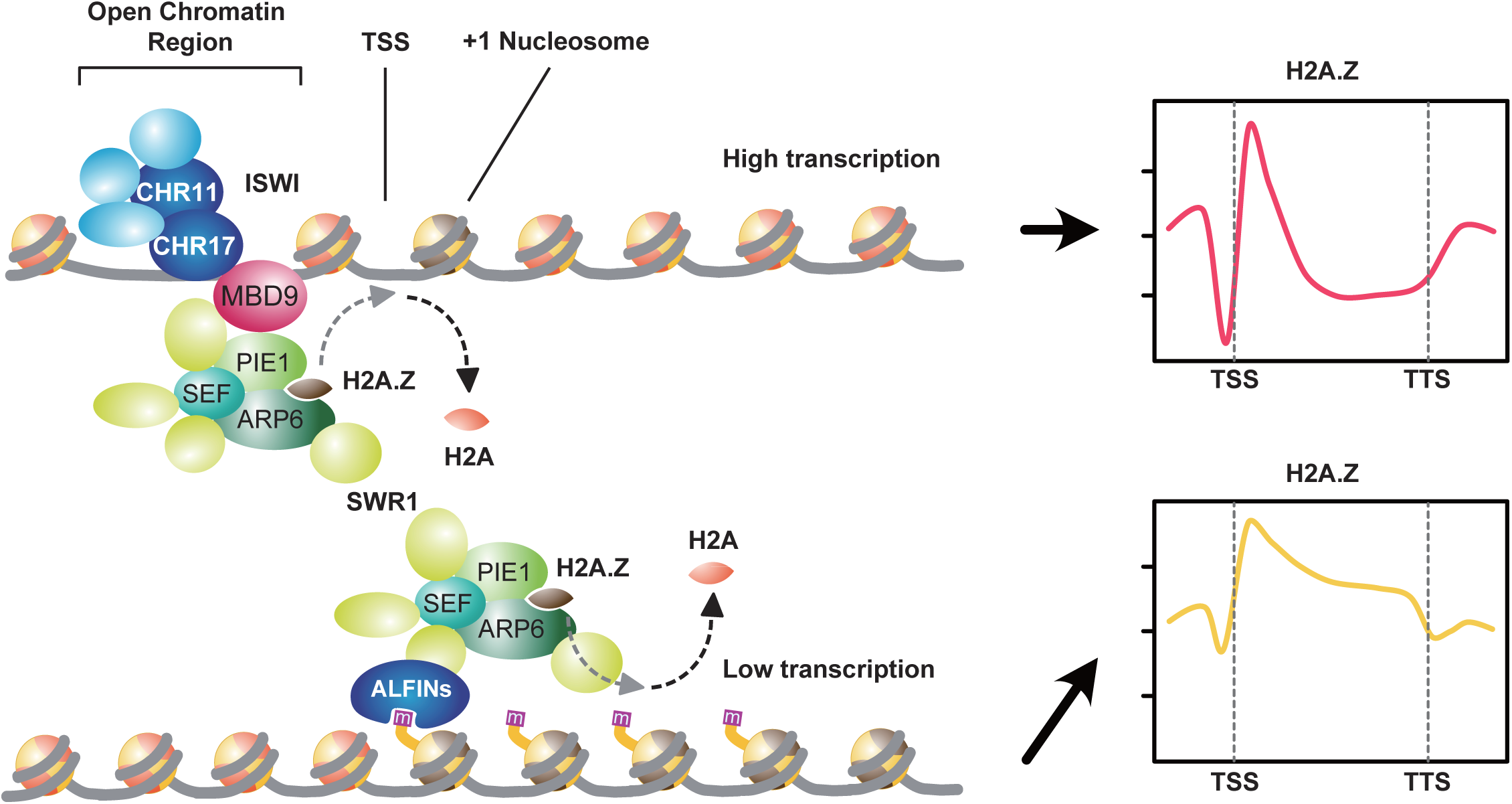
Proposed model of the role of ALFINs in transcription regulation. Unlike MBD9, which binds to the open chromatin regions and recruits the SWR1 complex to facilitate the deposition of H2A.Z at the +1 nucleosome, thereby enhancing transcription, ALFINs recognize H3K4me3 marks and recruit the SWR1 complex to deposit H2A.Z within the gene bodies of low-expressed genes.

## Discussion

The histone variant H2A.Z has been involved in many central aspects of plant biology (7). However, despite its importance, the precise mechanism by which H2A.Z is selectively deposited in some genomic regions is still unclear. Recently, MBD9 has been reported as a SWR1 interactor (18–20). MBD9 is likely responsible for recruiting SWR1 to the low-nucleosome regions immediately upstream of the TSS present in highly transcribed genes (19). This work focused on studying another SWR1 interactor, the AL proteins.

AL proteins have been previously described as H3K4me3 binding proteins using recombinant proteins and H3-modified peptides (30, 32). Our research has confirmed that AL5 binds H3K4me3 *in planta*. Moreover, AL5 and H3K4me3 strongly co-localize genome-wide. Importantly, however, the lack of functional AL has little to no effect on the accumulation of H3K4me3, indicating that ALs do not function in H3K4me3 dynamic regulation but rather act as H3K4me3 readers. In contrast, ALs are required for H2A.Z deposition/maintenance, as we observed a significant decrease in this variant at AL5-bound loci. This is consistent with previous reports of a physical interaction between several AL proteins and ARP6, a core component of the SWR1 complex (18–20), and suggests that AL proteins play a role in recruiting the SWR1 complex to a subset of H3K4me3 enriched loci.

We studied the phenotype of plants lacking regular expression of all the ALFIN-LIKE homologs found in Arabidopsis (*al7m*). These plants display severe abnormalities throughout development, indicating a vital role for these proteins under standard growing conditions. Moreover, we discovered that the majority of genes misregulated in the *al7m* typically exhibit little or no expression, and upon loss of ALs, their expression is significantly increased. We have also found that this subset of AL misregulated genes bears a combination of epigenetic features that are unusual for lowly transcribed genes, such as high levels of H3K4me3 and H3Ac - which typically decorate more highly transcribed genes - together with high levels of H3K27me3 and H2A.Z over the gene body - marks usually accompanying genes with low transcription.

Traditionally, H3K4me3 and H3K27me3 have been considered gene activation or suppression indicators, respectively. These two epigenetic marks have been long considered to be mutually exclusive. However, recent studies have revealed that they can appear together in specific genomic regions, resulting in a unique chromatin state (13, 14). The co-existence of H3K4me3 and H3K27me3 on the same gene is often called bivalency. Bivalency was first described in ESCs for cases where these two epigenetic marks were often found together in developmental genes. In differentiated cells, this bivalency was later resolved in favor of one of the marks, and only a handful of genes remained bivalent (45–48). In plants, bivalency is also present in a few developmental genes (49) and is also found in numerous stress-responsive genes (50–56), although it is worth noting that in these examples, direct evidence for the co-occurrence of H3K27me3 and H3K4me3 on the same (or adjacent) nucleosomes is often lacking. Interestingly, in *arp6* mutants, genes involved in biotic (57) and abiotic (58, 59) stress responses appear constitutively active in the absence of the triggering signal. Moreover, H2A.Z has also been described as a histone variant promoting gene expression responsiveness in Arabidopsis (9). In contrast, gene body methylation is anti-correlated with gene responsiveness and H2A.Z (60), although whether this is a direct effect or a consequence of low H2A.Z is still unknown. H3K4me3 is also associated with stress responsiveness and transcriptional memory (55, 56, 61, 62). Thus, it is tempting to speculate that AL proteins are specialized to regulate stress-responsive loci, directing the deposition of H2A.Z to genes bearing bivalent marks. In the future, it will be important to determine the nature and positioning of ALs within SWR1 to afford dynamic recruitment of the complex and to understand how the ALs themselves are regulated.

## Materials and Methods

### Plant material

All the plant materials used in this work are in Columbia (Col-0) background. We obtained *ALs* mutant alleles from ABRC (https://abrc.osu.edu): *al1-2* (Sail_1146_C08), *al2-1* (Sail_663_E04), *al3-1* (SALK_080056), *al5-1* (SALK_075676), *al6-1* (SALK_060877), and *al7-1* (SALK_032503). We generated *al4-2* using CRISPR-Cas9.

### Generation of al4-2

The sequence 5’-GAAACACAGTGTGGAGCAATG-3’ was used as the PAM cloned into the plasmid pHEE401(63). We used the pHEE401 plasmid containing our PAM in Arabidopsis plants. We then selected T_1_ transformants using MS plates containing 35mg/L of hygromycin. We screen the transformants by sequencing around the region targeted by our PAM. We harvested seeds from plants carrying mutations for further analysis in T_2_. In the T_2_, we selected plants negative for Cas9 and homozygous for the *al4-2* mutation. We backcrossed the original *al4-2* twice against Columbia to minimize possible off-target mutations.

### RNA-Seq

We extracted total RNA from 28-day-old plants using TRIzol (Invitrogen). The supernatant containing total RNA was further purified using AxyPrep Multisource Total RNA (Axygen). We sent RNA samples from five biological replicates to BGI genomics (https://www.bgi.com) for sequencing.

The raw reads were first filtered with fastp (version 0.22.0) for low-quality reads, then mapped against the reference genome (TAIR10) using hisat2 (version 5.4.0). The transcriptome was quantified using a subread software, and the GTF file annotated by araport11 was used as the annotation file. We used DEseq2 to analyze differential expression. We used the PCAtools software package to perform PCA analysis and tidyverse to process the data further. We used ggplot2 to draw the volcano plot, boxplot, and pie chart. We used ComplexHeatmap to draw heat maps and classify genes according to expression and correlation coefficient. We used ClusterProfiler for the GO enrichment analysis of differentially expressed genes. we used a q-value cut off=0.05, and a p-value cut off=0.05. We used org.at.tail.db software package for GO annotation.

### Chromatin immunoprecipitation

For the ChIPs of the *al7m*, 0.5g 40-d-old seedlings were harvested alongside Col-0 control. For the ChIPs of AL5-FLAG, 2.0g 15-d-old whole plants were harvested from ½ MS plates alongside Col-0 control. ChIPs were performed as previously described(64) https://doi.org/10.7554/eLife.89353.2. For all ChIPs, two biological replicates were performed. The antibodies used were: anti-FLAG M2 (F1804, Sigma-Aldrich), anti-H3K4me3 (ab8580, Abcam), anti-H2A.Z (AS10718, Agrisera) and anti-H3 (ab1791, Abcam). Libraries were generated with NuGen Ovation Ultra Low System V2 kits following the manufacturer’s instructions. The libraries were sequenced for PE150 reads in Illumina NovaSeq 6000.

The reads were aligned to the TAIR10 reference genome using Bowtie2 (version 2.5.3), the duplicated reads were removed by Samtools (version 1.19.2). Deeptools (version 3.5.5) was used to generate the tracks using RPGC for normalization. As there were few differences between the biological replicates in all ChIPs, we merged the biological replicates for downstream analysis. The IGV genome browser was used to visualize the data. MACS2 (version 2.2.9.1) was used for peak calling and differential peaks were identified by DiffBind in R. Peaks were annotated by ChIPseeker in R. Deeptools was used to generate the metaplots and heatmaps.

### Co-immunoprecipitation

Jin-Song Zhang and Shou-Yi Chen from the State Key Laboratory of Plant Genomics, Institute of Genetics and Developmental Biology, Chinese Academy of Sciences generously provided the α-AL5 antibody(38). We used the Dynabeads antibody coupling kit (Invitrogen) to attach the α-AL5 antibody to magnetic beads. We started by grinding 1 gram of floral tissue in liquid nitrogen. Then we added 10 mL of IP buffer (50 mM Tris pH 7.6, 150 mM NaCl, 5 mM MgCl2, 10% glycerol, 0.1% NP-40, 0.5 mM DTT, 1 mM, β-mercaptoethanol, 1 mM PMSF, and 1xProtease Inhibitor cocktail (MCE)). We incubated the mixture on ice for 15 minutes and then spun it at 4,000g for 10 minutes at 4°C. We then filtered the supernatant through a double Miracloth layer (Millipore). We added the magnetic beads previously coupled to the α-AL5 antibody and incubated them for 1 hour at 4°C with gentle rotation. We washed the beads five times with IP buffer and then released the immunoprecipitate by boiling. Finally, we performed Western blotting and detection using Streptavidin-HRP (1:5,000) (Cell Signaling Technology) or α-H3K4me3(1;2,000) (Millipore) antibody.

### Protein extraction and Western blotting

We conducted an analysis of protein expression using western blots. To detect non-histone proteins, we first ground approximately 100mg of fresh tissue in liquid nitrogen and added 250μL of protein extraction buffer (50mM Tris HCl, pH-7.4, 1% β-mercaptoethanol, 12% sucrose, 0.1% Triton X-100, 5mM PMSF). We shook the mix vigorously and let it sit on ice for 15 minutes before spinning it at 12,000g at 4°C. We then transferred the supernatant to a fresh tube, repeating this step twice. Before loading, we mixed the samples with an appropriate volume of 4x protein loading buffer (250mM Tris-HCl pH = 6.8, 400mM DTT, 8% SDS, 40% glycerol, 0.1% bromophenol blue). We denatured the proteins by boiling them for 15 minutes. We used the EpiQuik^TM^ total histone extraction kit (EPIGENTEK) to detect histones and followed the manufacturer’s instructions. We loaded 10-20μL of proteins into 4–12% for total protein or 12% acrylamide gels for histone extracts (Invitrogen). We transferred the proteins from the gel to an Immobilon-P membrane (Millipore) at 400mA for 2 hours. The antibody dilution used was as follows: Streptavidin-HRP (1:5,000) (Cell Signaling Technology) or α-H3K4me3 (1:2,000) (Millipore) antibody. According to the manufacturer’s recommendations, we detected the HRP signal using the Super Signal West Pico Plus^TM^ Chemiluminescent Substrate kit (Thermo).

## Funding

The Ausin lab work was supported by the National Science Foundation of China (NSFC) 321703059 and 32370580 grants. The work in the Harris la was supported by the Royal Society (URF\R1\201016) and a European Research Council (ERC) / UK Research and Innovation (UKRI) grant (EP/X025306/1).

## Author contribution

LHX, YFW, XYL, QH, VA, and IA conducted the experiments, LHX and JJX analyzed genomic data, CJH, YFW, and IA designed the experiments, and CJH and IA wrote the manuscript with the participation of all the other authors.

## Acknowledgments

We thank Jin-Song Zhang and Shou-Yi Chen for the kind gift of the α-AL5 antibody and Jiang Dan for her technical assistance.

## Declaration of interest

The authors declare no competing interest.

## Data availability

All sequencing data are accessible at the NCBI Gene Expression Omnibus (GEO) under accession number GSE270861 (ChIP-seq) and GSE270862 (RNA-seq).

---

**Supplemental Figure 1.**
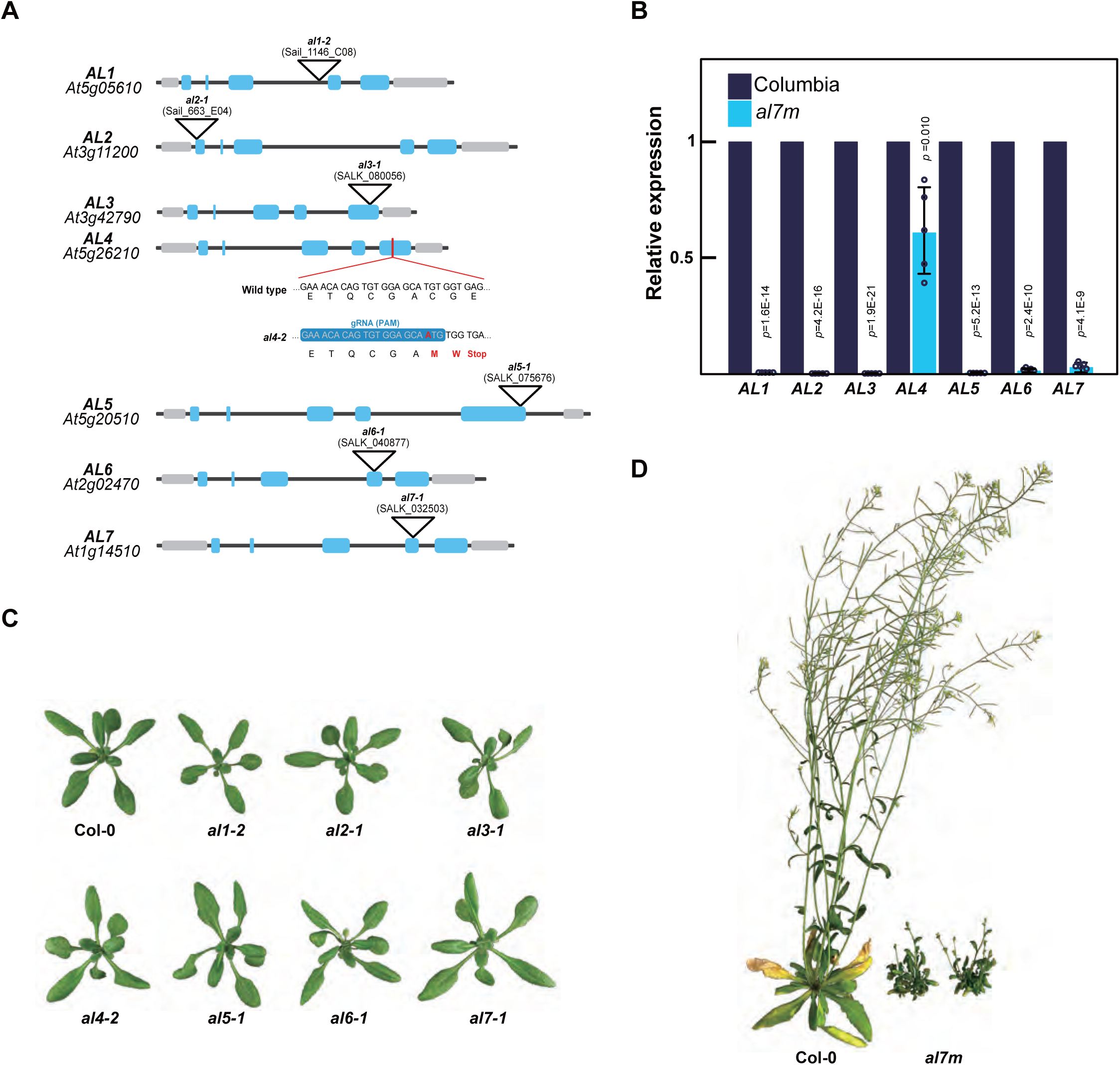
alfin-like single mutants’ alleles and alfin-like mutants’ phenotype. (A) graphical representation of the AL family genes showing the nature and position of the mutations carried by the alleles used in this work. Inverted triangles represent T-DNA insertions, and the red line represents appoint mutation. (B) RT-qPCR data showing the expression levels of the AL genes in the al7m background and wildtype. Data are presented as the average of five biological replicates. Error bars represent SD, and significance was calculated using Student’s T-Test. (C) The visual phenotype of the wildtype and alfin-like single mutants. (D) The visual phenotype of the wildtype and al7m plants at late growth stage.

**Supplemental Figure 2.**
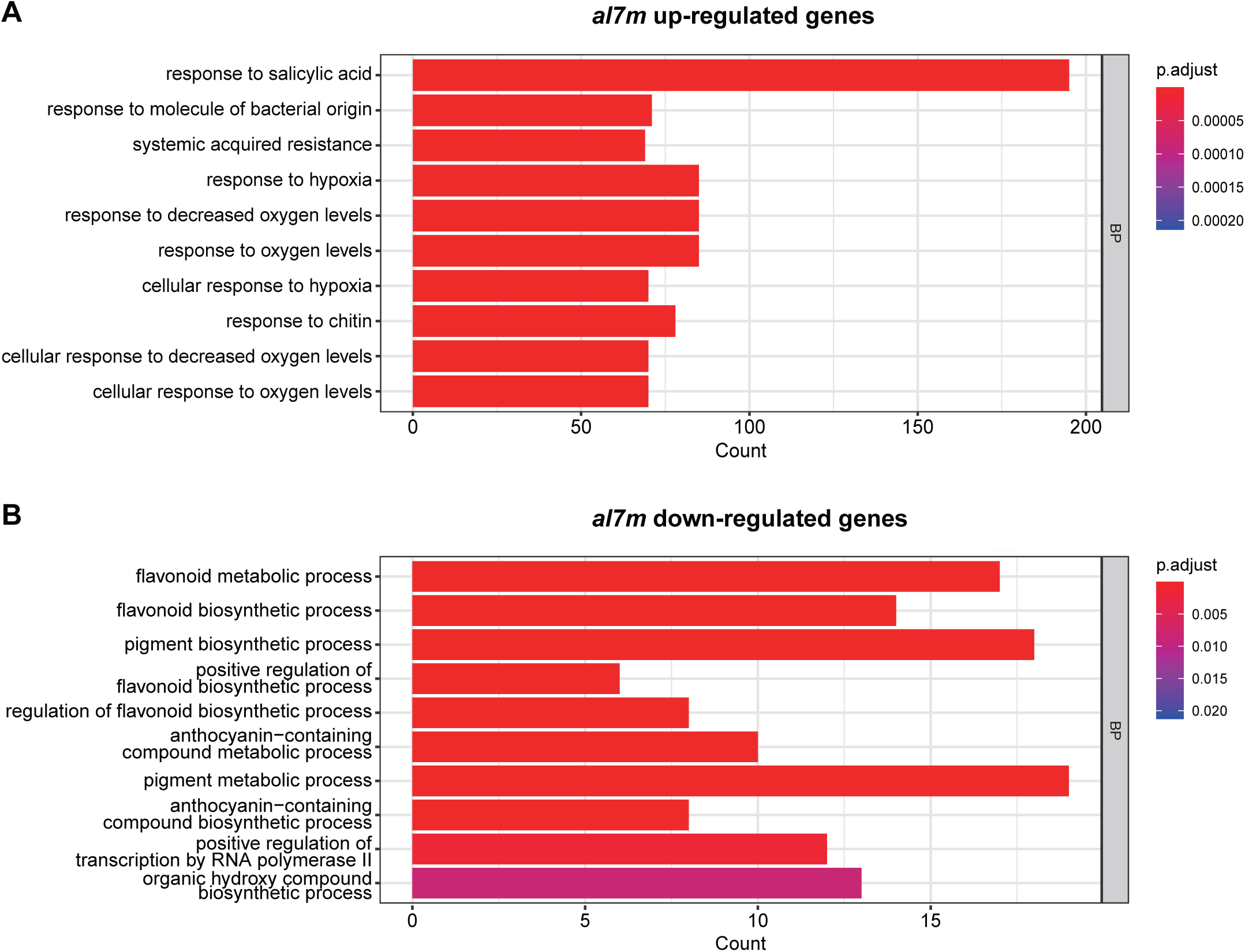
Gene ontology analysis (biological processes) of al7m up-regulated genes (A) and al7m down-regulated genes (B).

**Supplemental Figure 3.**
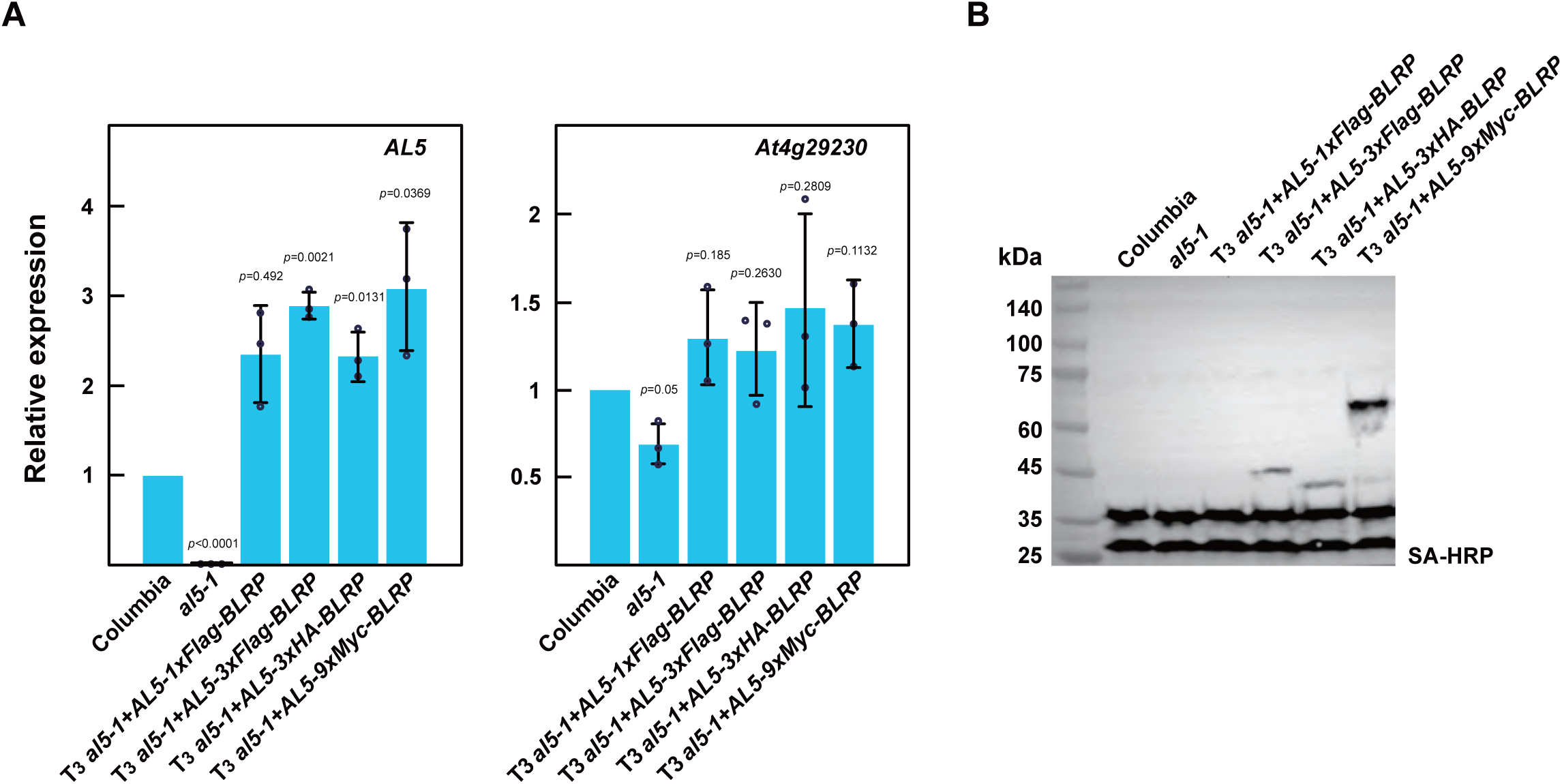
Complementation of the alfin-like5 mutant. (A) RT-qPCR showing the expression levels of the AL5 and At4g29230 genes in Columbia, al5-1, and several AL5-tagged complementing lines. Error bars represent SD, and statistical significance was calculated using Student’s T-Test. (B) Western blot hybridization showing the expression of the AL5-tagged complementing lines at the protein level. HRP-labelled streptavidin was used for protein detection.

**Supplemental Figure 4.**
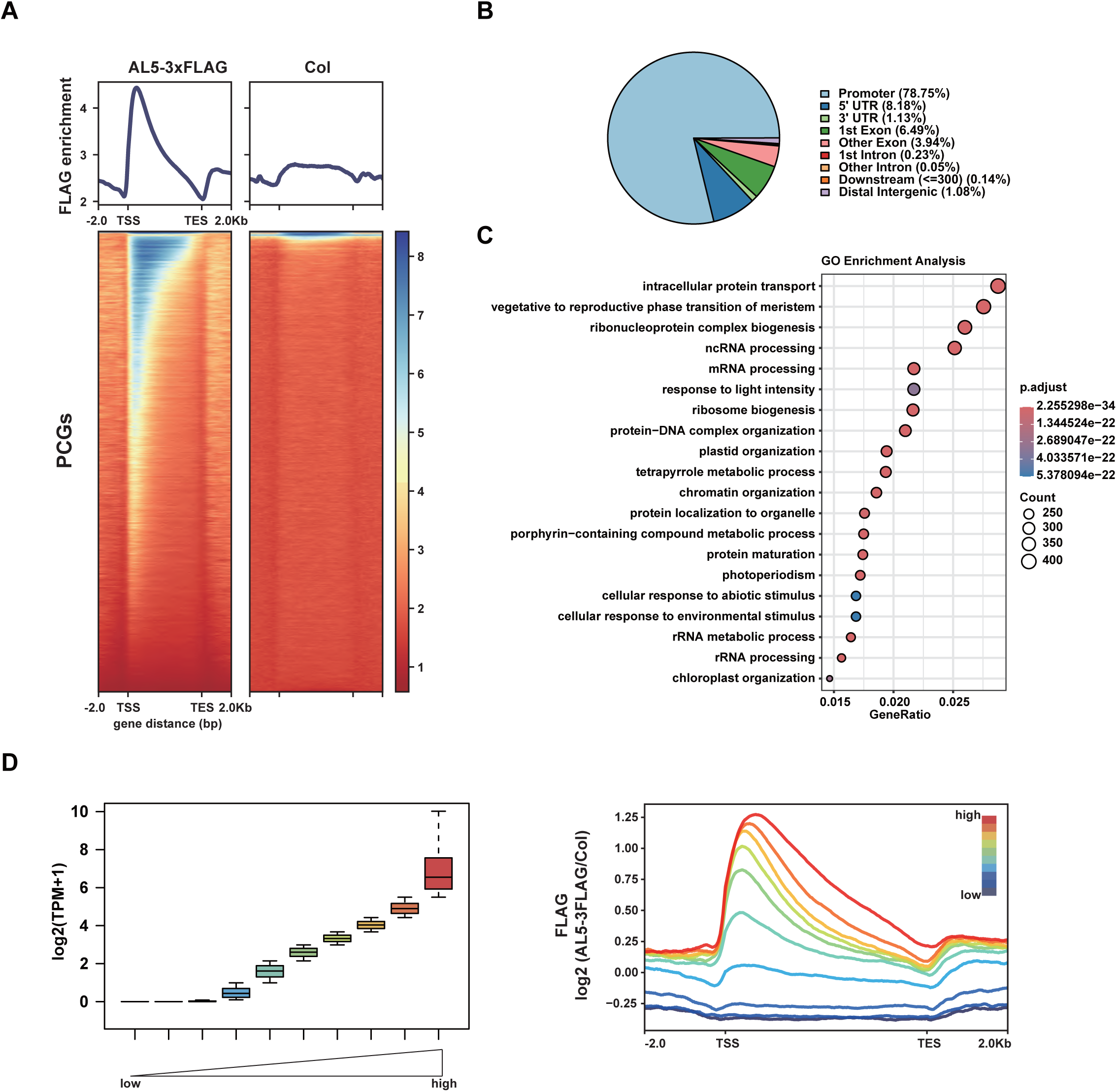
AL5-FLAG ChIP-seq validation. (A) Metaplots and heatmaps showing the ChIP-seq signals of FLAG over PCGs in AL5-3Xflag and wildtype. (B) Pie Chart showing the annotations of the AL5-FLAG binding genes (n=14866). (C) Gene ontology analysis (biological processes) of AL5-FLAG binding genes. (D) Boxplot and metaplot showing sanity check of genes expression levels (left) and average occupancy of AL5 over genes with different expression levels (right).

**Supplemental Figure 5.**
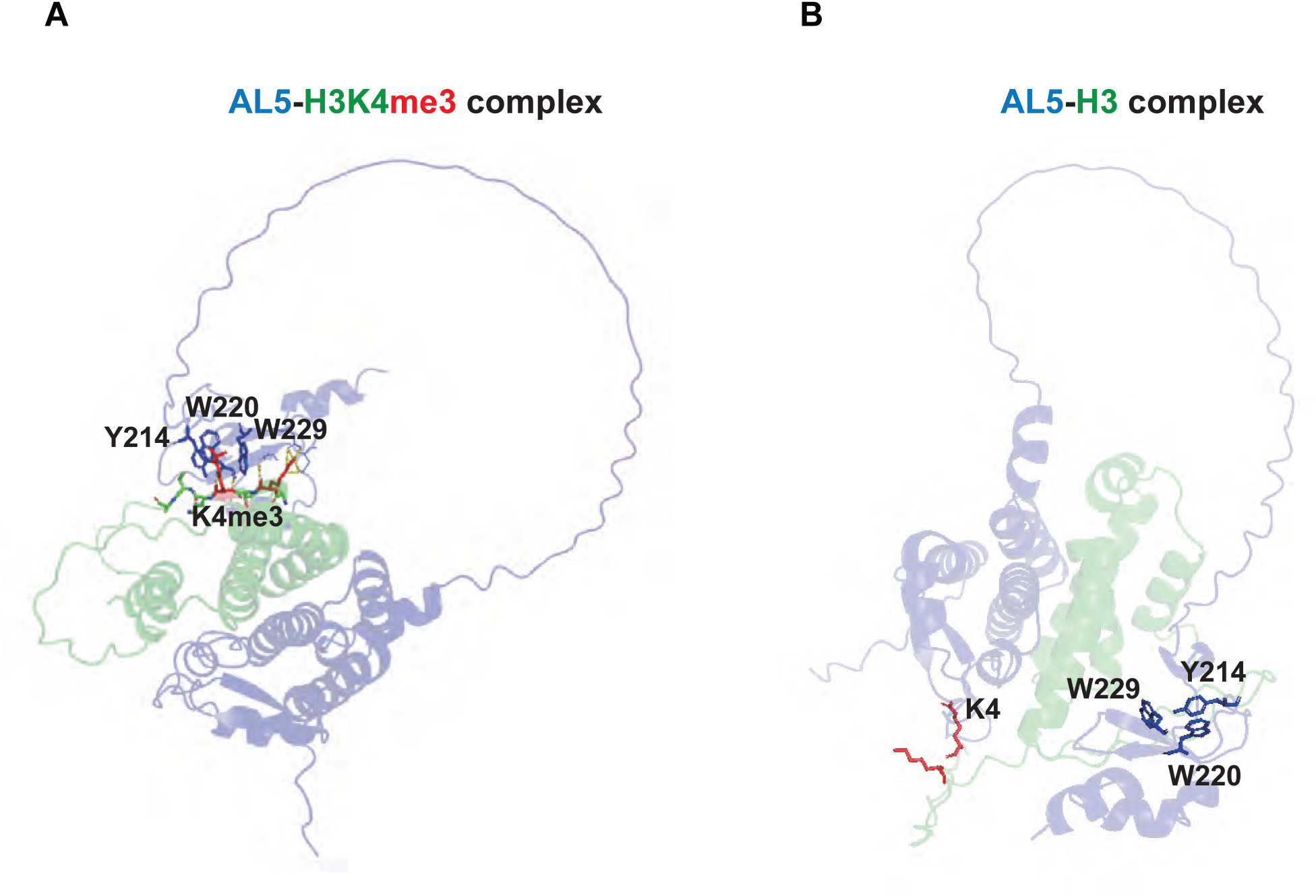
AlphaFold 3 predicted structure models of AL5-H3K4me3 complex (A) and AL5-H3 complex (B). The AL5 is shown in blue, H3 is shown in green and the histone H3K4me3 is labeled. AL5 residues involved in H3K4me3 recognition are labeled.

**Supplemental Figure 6.**
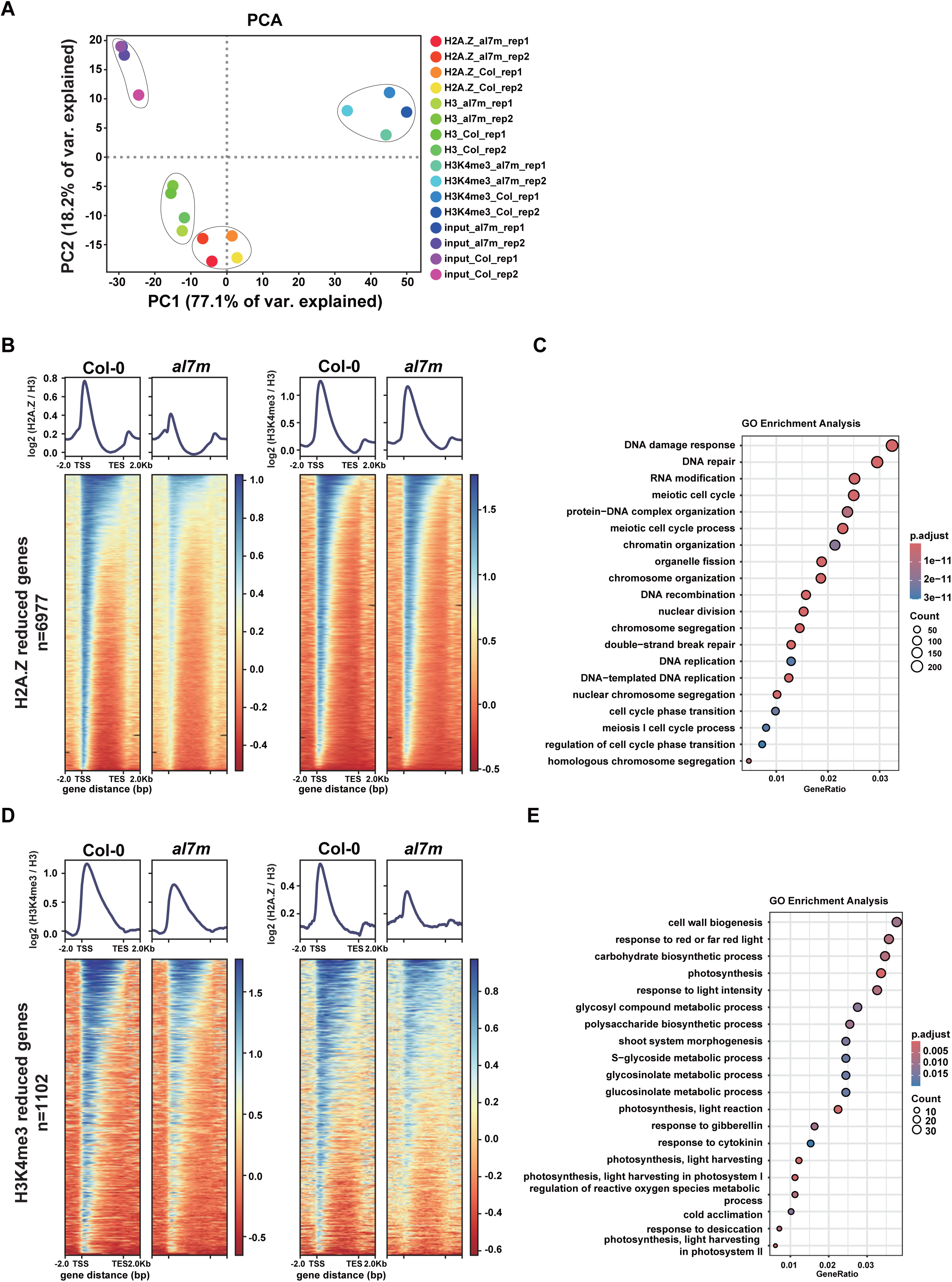
H3K4me3 and H2A.Z ChIP-seq validation. (A) PCA of ChIP-seq signals of H3K4me3, H2A.Z, H3, and input in Col-0 and al7m, each with two biological replicates. (B) Metaplots and heatmaps showing the ChIP-seq signals of H2A.Z and H3K4me3 over genes with reduced H2A.Z (n = 6977) in al7m and Col-0. (C) Gene ontology analysis (biological processes) of H2A.Z reduced genes. (D) Metaplots and heatmaps showing the ChIP-seq signals of H3K4me3 and H2A.Z over genes with reduced H3K4me3 (n = 1102) in al7m and Col-0. (E) Gene ontology analysis (biological processes) of H3K4me3 reduced genes.

**Supplemental Figure 7.**
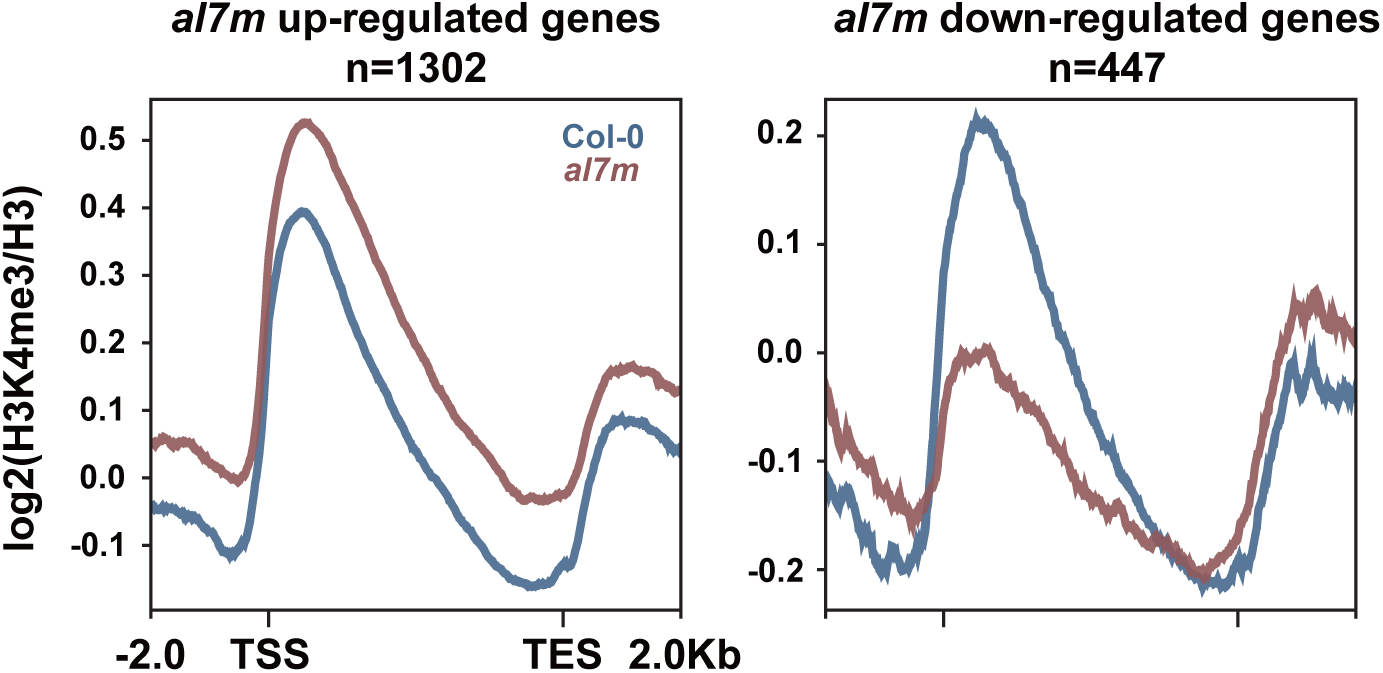
Metaplots showing the ChIP-seq signal of H3K4me3 in Col-0 and al7m over al7m deregulated genes.

**Supplemental Figure 8.**
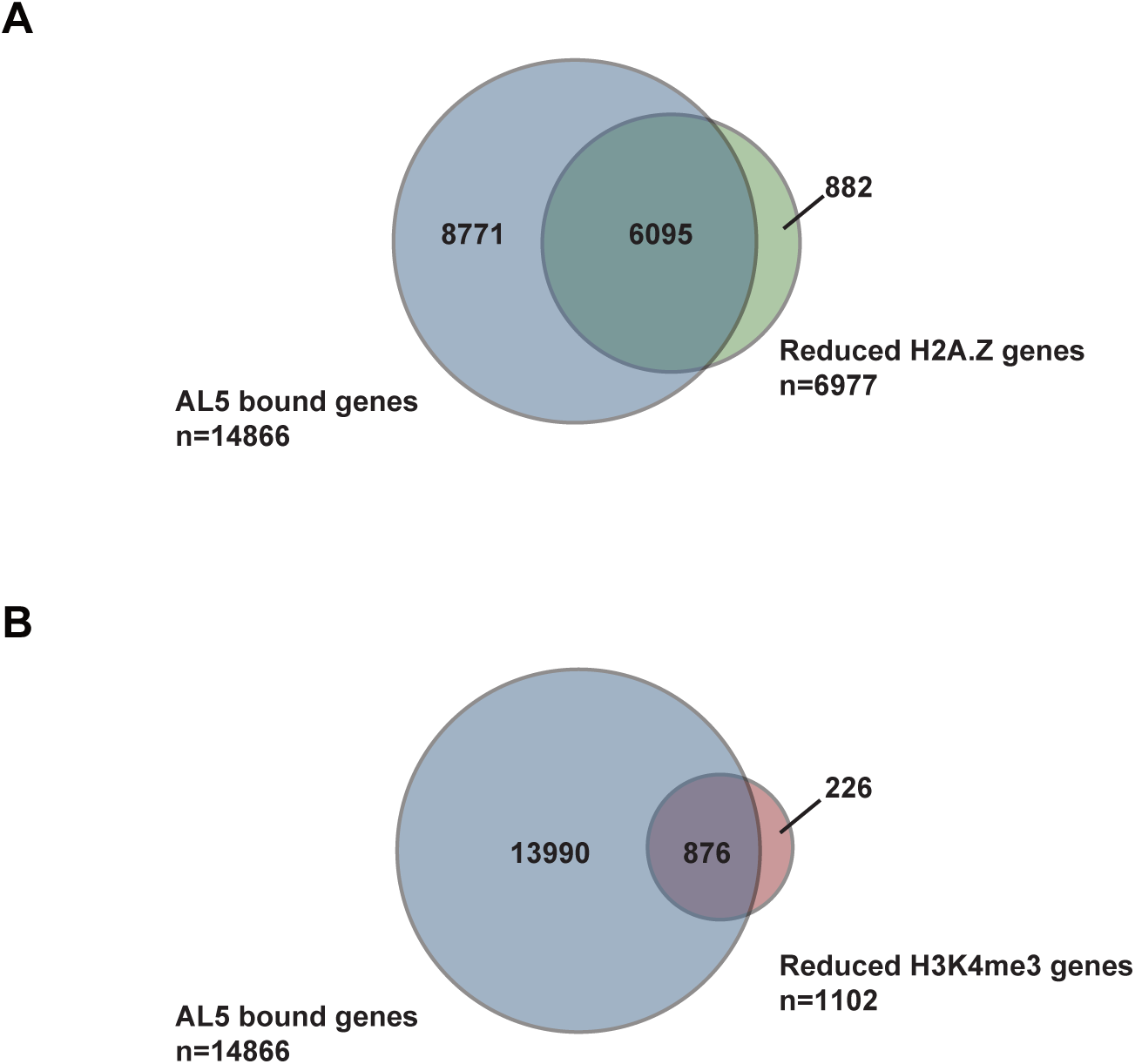
Venn diagram analysis showing the overlap of AL5 bound genes (n=14866) with reduced H2A.Z genes (A) and reduced H3K4me3 genes (B).

**Supplemental Table1.** Summary of AL5-9MYC IP Mass spec.

| Protein | Annotation | AL5-9Myc | Columbia |
| --- | --- | --- | --- |
| AT5G20510 | AL5 | 7 | 0 |
| AT3G12810 | PIE1 | 22 | 0 |
| AT5G22330 | RuvB-like protein 1 (EC 3.6.4.12) | 16 | 2 |
| AT5G67630 | RuvB1 | 13 | 0 |
| AT1G18450 | ARP4 | 10 | 0 |
| AT2G47210 | SWC4 | 8 | 0 |
| AT2G36740 | SWC2 | 7 | 0 |
| AT5G26210 | AL4 | 6 | 1 |
| AT2G02470 | AL6 | 5 | 2 |
| AT3G42790 | AL3 | 4 | 0 |
| AT3G33520 | ARP6 | 4 | 0 |
| AT3G18790 | MVE11_17 | 4 | 1 |
| AT4G35785 | RNA-binding (RRM/RBD/RNP motifs) family protein | 4 | 1 |
| AT2G29210 | Splicing factor PWI domain-containing protein | 4 | 1 |
| AT5G48880 | KAT5 | 4 | 1 |
| AT3G11200 | AL2 | 3 | 0 |
| AT4G19400 | Profilin family protein | 3 | 0 |
| AT1G14830 | Phragmoplastin DRP1C (EC 3.6.5.-) | 3 | 0 |
| AT2G34750 | RNA polymerase I specific transcription initiation factor RRN3 protein (Uncharacterized protein) | 3 | 0 |
| AT5G37055 | SEF | 2 | 0 |
| AT1G54250 | NRPB8A | 2 | 0 |
| AT3G20550 | DDL | 2 | 0 |
| AT1G10200 | WLIM1 | 1 | 0 |

## Notes

### Competing Interest Statement

The authors have declared no competing interest.

